# Hormones: What are they good for?^‡^

**DOI:** 10.64898/2026.08.24.746760

**Authors:** Sean A. Ridout, Priyathama Vellanki, Ilya Nemenman

## Abstract

Animals use long-range signals, such as hormones and neural signals, to coordinate the actions of distant organs. There is no precise, quantitative framework that explains the problems these control systems must solve and thus predicts their behavior under varied conditions. We consider this problem in the context of blood glucose regulation by the hormone insulin, the failure of which produces diabetes. We show that existing mathematical models of glucose regulation admit equivalent control strategies with no hormones at all, and thus cannot explain the need for hormonal regulation. We therefore introduce a minimal model of inter-organ variations in local glucose, and show that control strategies based on local glucose measurements face severe trade-offs between different control objectives. In contrast, we show that hormonal control signals from the pancreas can overcome these limitations. By exposing the benefits of hormonal control, our work paves the way to a detailed understanding of physiological design principles, with possible implications for the engineering of an artificial pancreas.

## I. INTRODUCTION

Hormonal signaling is one of the central features of animal physiology: virtually every aspect of metabolism, growth, reproduction, and development is regulated by hormones, from insulin and glucagon to cortisol, thyroid hormone, estrogen, and testosterone [1]. Yet, despite their importance and more than a century of experimental and clinical study, there is surprisingly little quantitative understanding of what hormones are *for* at the level of physiological design principles. Recent work has addressed design principles for individual ligands [2] and the regulatory circuit [3]. Neither addresses the spatial organization of the body, although hormones have been understood since their discovery to “coordinate” the actions of distant organs [4, 5]. Indeed, it has never been made precise what problems endocrine signaling solves that could not, in principle, be solved by *local* regulation within each organ or tissue. Why, then, does physiology rely so heavily on long-range chemical messengers?

We address this question in the context of human glucose homeostasis, which is regulated primarily by the hormone insulin [1]. Glucose is a vital fuel, and its deficiency in the brain is acutely life-threatening [6]. In excess, however, it is a driver of long-term pathology, most notably diabetes, one of the most prevalent and costly chronic diseases worldwide [7]. Its concentration in the blood is therefore tightly regulated by the action of several organs, including the liver (which can either release or store glucose) and muscle and adipose tissue (which consume it or release substrates for its production) [1]. This regulatory system has been studied in great detail, due to its clinical importance. Still, it remains unclear why glucose control should require a circulating hormonal signal when each organ could regulate its own glucose uptake or production based only on the local glucose concentration it experiences. In the bodies that evolution has given us, removing insulin results in diabetes. But could evolution have instead chosen a different glucose control system without endocrine signaling?

One reason this question is unanswered is that theoretical work on glucose regulation has been driven by the features of diabetes, and therefore frames the problem as restoring global blood glucose to a fixed set point [8– 12]. This perspective is useful in clinical applications, but leaves the design goals of the intact system unexamined. Conceptually, this focus is questionable: for most of evolutionary history, glucose was primarily a fuel rather than a poison, and ensuring its reliable delivery to tissues was a more important challenge than rapidly metabolizing a can of Coke. Moreover, the many complexities of the insulin–glucose system seem unnecessary to merely return glucose to a set point. For example, insulin secretion depends not only on glucose concentration but also on its rate of change [13, 14], and is pulsatile even in steady conditions [15]. Further, glucose homeostasis also relies on the action of multiple other hormones [1, 16–18].

Here we propose that the necessity of hormonal signaling in this system follows from a different control objective than the usually assumed restoration of a precise global set point. We hypothesize that its primary goal is *to ensure robust and reliable delivery of glucose to multiple organs with heterogeneous, time-varying demands*. Thus, glucose regulation is a whole-body, dynamic resource-allocation problem, requiring coordination between spatially separated sites of production, storage, and consumption. The anticipatory allocation of fuel to organs according to predicted demand is known in the physiology literature as *allostasis* [19, 20]. Like the allostasis concept, our analysis is built around the view that meeting the body’s multiple competing demands, rather than merely maintaining a set point value, is the ultimate goal of physiological regulation. Here, however, we treat the spatial aspects of this problem, and leave anticipation of future demands for future work.

Specifically, we show that in a well-mixed system, any hormonal control strategy could, in principle, be replaced by an equivalent local control circuit, making hormonal control merely one solution among many. We then build a minimal model of glucose production and consumption in spatially separated organs, parametrized by existing data. Using this model, we show that, in such a distributed system, local control cannot adequately meet heterogeneous, time-varying demands. Finally, we show how hormonal control can surpass these limits, in agreement with known physiology. Thus, hormonal glucose control achieves crucial objectives that cannot be met by local control.

In the context of glucose regulation and diabetes, understanding what the hormonal regulatory network is designed to achieve—and thus why it is organized the way it is—should help predict failure modes and guide therapeutic interventions. More broadly, we provide a quantitative approach to understanding when hormonal signaling is not just one solution among many, but a *necessary* tool for coordinating spatially distributed physiological processes. We thus see that the ubiquity of hormones across animal physiology is due not just to historical contingency, but to biophysical and physiological constraints.

## II. RESULTS

### A. Without glucose gradients, hormones are dispensable

A standard explanation of the role of insulin is that it ensures non-obligate users of glucose spare glucose for the brain, drawing on blood glucose primarily when it is in excess [21]. It seems, however, that evolution could have *instead* achieved this goal using nonlinear, tissue-specific responses to glucose itself, without requiring a hormonal signal.

To formalize this point, consider a standard model [8] of the feedback loop between the blood glucose concentration at time *t, G*(*t*), and the hormone insulin, *I*(*t*):

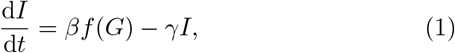

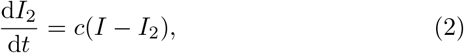

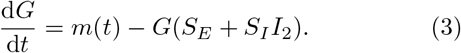

Here *I*_2_(*t*) is the insulin in the “remote compartment,” such as muscles, where it acts on tissues. *f* (*G*) is a (nonlinear) secretion function. The constants *β, γ*, and *c* are the maximum insulin secretion rate, the insulin clearance rate, and the exchange rate of insulin between the compartments, respectively. *m*(*t*) represents glucose input (e.g., meals and hepatic output), while *S*_*I*_ and *S*_*E*_ set insulin-dependent and insulin-independent glucose clearance rates. *γ*^−1^ and *c*^−1^ are typically minutes to tens of minutes. Crucially, in this model glucose forms a single, well-mixed pool *G*(*t*). The model lacks a description of how glucose gradients might exist between organs, and each organ might sense and respond to a different glucose concentration, affecting glucose dynamics.

Equations (1) and (2) can be formally solved, showing that insulin encodes a nonlinearly filtered *history* of recent glucose concentrations. Assuming the system has run long enough for transients to decay (*SI Appendix*, Section S1),

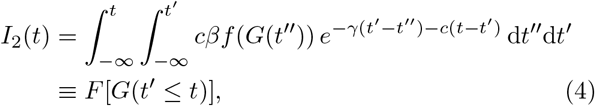

which defines a functional *F* of the past glucose trajectory.

Therefore, this hormone-mediated regulation, Eqs. (1)– (3), is mathematically equivalent to a hormoneless control algorithm, in which tissues respond directly to glucose as

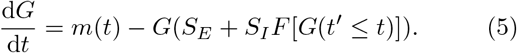

Thus, in this model insulin can be “integrated out,” leaving an equivalent local response with short-term memory. Because every tissue is exposed to the same *G*(*t*), each can reconstruct its own filter *F* [*G*] from the glucose it senses locally, so the circulating insulin signal conveys nothing a tissue could not compute on its own. Such a response could be implemented using intracellular signaling dynamics, which can perform temporal filtering of time-varying chemical inputs [22], without requiring any endocrine signal. Even benefits of β-cell adaptation, such as “dynamical compensation” [11], could be produced by adaptation of a coefficient inside the functional *F* .

This observation is not unique to this model, or even to glucose regulation. For example, a model of interactions between insulin and glucagon [23] and a model of hormonal control of blood calcium [24] can be rewritten as hormoneless models in this way (*SI Appendix*, Section S2). In more sophisticated compartmental models of insulin action [25], some aspects cannot be rewritten this way, but the key ones can (*SI Appendix*, Section S2 D).

Thus, if glucose is well-mixed, hormonal control buys nothing that a purely local response could not, and such models cannot explain why evolution did not dispense with hormones altogether. We therefore hypothesize that insulin’s role is to coordinate glucose regulation across the *glucose gradients* between organs with heterogeneous, time-varying demands.

### B. A minimal distributed glucose regulation model

To show that distributed physiological dynamics pose control problems that are absent in models with a single, well-mixed glucose pool, and thereby require hormones, we introduce a minimal model that explicitly includes *spatial variation* in metabolite and hormone concentrations.

Blood leaves the heart via the aorta and is distributed to organs by arteries and arterioles. Each organ is fed by capillaries, then drained by venules and veins. We begin with known anatomy [26], coarse-grain this branching structure, and neglect organs and vessels that are not needed in our analyses (e.g., the pulmonary and coronary circulations), yielding the network shown in Fig. 1.

**FIG. 1.**
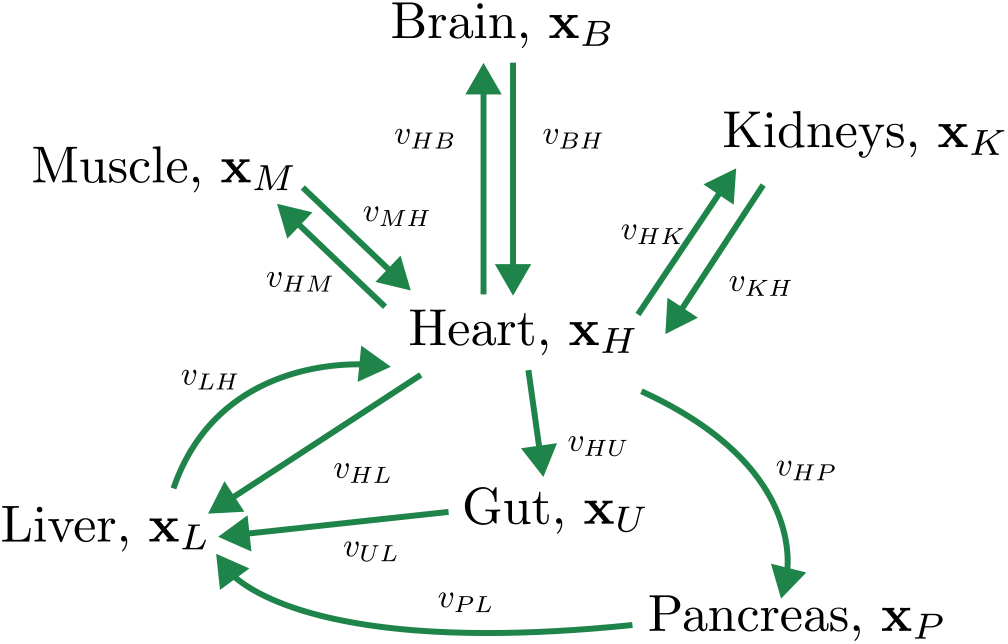
Our minimal model of distributed control problems: Each organ *i* has a vector of metabolite and hormone concentrations **x**_*i*_. For example, *G*_*L*_ is the glucose concentration in liver capillary blood. Organs can only directly respond to or adjust the metabolite and hormone concentrations in their own capillary beds. Flows with volumetric rates *v*_*ij*_ and time delays *τ*_*ij*_ transport chemicals from organ *i* to organ *j*.

We make two further simplifying approximations. First, we treat the blood in the capillary bed of each organ *i*, of volume *V*_*i*_, as well mixed. We thus assign concentrations *x*_*i*_ to a molecule *x* in organ *i*, and **x**_*i*_ refers to all species simultaneously. Second, we model transport between organs as plug flow, neglecting dispersion (see *SI Appendix*, Section S3 A). Thus, blood moves from organ *i* to organ *j* with volumetric flow rate *v*_*ij*_ and time delay *τ*_*ij*_.

In each organ, species are produced or consumed at rates determined by the local concentrations and conditions, with a net production rate **g**_*i*_(**x**_*i*_, *t*). The dynamics thus obey a system of delay differential equations (DDEs),

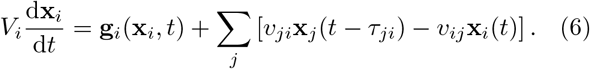

We will usually analyze a representative set of adult resting flow rates *v*_*ij*_ (*SI Appendix*, Section S3 B, Table S1). In reality, flow rates vary (e.g., changing in exercise to meet oxygen demands), and we will discuss this variation when relevant. Parameters also vary from person to person, making control *harder* ; we leave this issue for future work.

We will often need the glucose component of **g**_*i*_(**x**_*i*_, *t*), which we call *m*_*i*_(**x**_*i*_, *t*). Throughout, organ subscripts denote the heart (*H*), brain (*B*), muscle (*M*), liver (*L*), kidneys (*K*), gut (*U*), and pancreas (*P*). Where it does not cause confusion, we drop the explicit time dependence and write *m*_*i*_(**x**_*i*_). *R*_0_ *>* 0 denotes the total rate of glucose utilization and production at fasting steady state.

This distributed description is the minimal model in which different organs can experience different concentrations at the same time and are coupled only through transport with finite delays. This spatial separation breaks the equivalence between hormonal and local control that we derived above. We now use this model to analyze three physiologically relevant conditions: fasting, exercise, and a carbohydrate meal. We will show that hormoneless control faces severe trade-offs between different control objectives, which are eliminated by hormonal control.

### C. Spatial variations of glucose during fasting

We begin by analyzing fasting conditions, showing that spatial gradients in our model are large enough to matter and match experimental values. At steady state, our distributed model implies that an arteriovenous (AV) glucose concentration difference is proportional to tissue uptake or production, see Eq. (6). These differences arise from mass balance and are independent of the body’s control algorithm.

After an overnight fast, total endogenous glucose production *R*_0_ in a 70 kg adult man is roughly 140 mg*/*min, with ∼80% produced in the liver and ∼20% in the kidneys [27]. Of this, ∼50% is consumed by the brain, ∼10% by the gut, liver, and nearby tissues, ∼20% by muscle, ∼10% by blood and skin, ∼5% by the kidneys, and ∼5% by adipose tissue [27], giving estimates of *m*_*i*_. With our sign convention (*m*_*i*_ *>* 0 for net release to blood), and using estimates for the resting flow rates from Table S1, this results in the steady-state balance between production/consumption and inflow–outflow

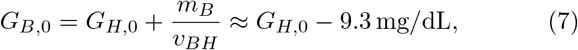

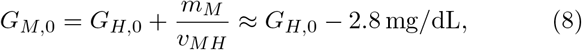

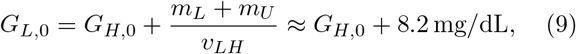

where the subscript 0 denotes overnight fast condition.

Thus, a substantial gradient exists between the main site of glucose consumption (the brain) and the main site of glucose production (the liver):

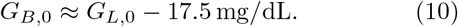

These *G*_*i*,0_ represent capillary concentrations within each organ. Since glucose diffuses down concentration gradients, differences between intratissue or intracellular concentrations are larger. We also note that these estimates are comparable to measured arteriovenous differences in healthy fasting subjects (∼10 mg*/*dL across the brain [28] and ∼2 mg*/*dL across forearm muscle [29]).

### D. Hormoneless instantaneous control

We now analyze a broad class of hormoneless control strategies in three conditions: exercise, meals, and fluctuations in blood flow or demand. We consider *hormoneless instantaneous local control* : each organ *i* sets its net glucose flux according to a function *m*_*i*_(*G*_*i*_(*t*)) of its own current capillary glucose concentration, with no access to circulating signals, or to the history of glucose or of its own activity (we will return to algorithms using history at the end). Each function *m*_*i*_(*G*_*i*_) is different, reflecting organ-to-organ differences in cellular physiology (e.g., different glucose kinases [30]). While in real organisms some failures of such control might be mitigated by neural signaling, our goal here is to isolate the limitations of purely local, instantaneous regulation.

Under all three conditions, we will show that these control algorithms face unavoidable physiological trade-offs. The origin of these trade-offs is that, as seen in Eqs. (7)–(9), glucose gradients are large, and change in response to changes in glucose production and consumption— including the very changes the body uses to achieve control. Thus, the effects of organs on glucose contaminate their ability to infer glucose demands and levels elsewhere in the body, which is needed to coordinate whole-body regulation. The next three sections present these analyses.

#### 1. Hormoneless instantaneous control cannot prevent brain hypoglycemia during exercise

We first show that a hormoneless instantaneous control algorithm cannot both keep brain glucose safe during exercise and maintain normal fasting glucose. This limitation holds even under optimistic assumptions about how muscle energy demand is met and is independent of the details of the control laws *m*_*i*_(*G*_*i*_).

The liver can either release glucose into the blood or store it as glycogen, so its net flux *m*_*L*_(*G*_*L*_) can be positive or negative. In contrast, in humans, skeletal muscle does not release glucose, so *m*_*M*_ (*G*_*M*_) ≤ 0. The brain is a largely obligate consumer, with uptake *m*_*B*_ ≈ const *<* 0 set by its metabolic needs, and the kidneys produce glucose during fasting, with net rate *m*_*K*_ *>* 0. We define the fasting steady-state rates *R*_*M*,0_ ≡ − *m*_*M*_ (*G*_*M*,0_) and *m*_*L*,0_ ≡ −*m*_*L*_(*G*_*L*,0_), and similarly for other organs.

All of the local control functions *m*_*i*_(*G*_*i*_) must be monotonically decreasing: e.g., high *G*_*L*_ should tell the liver to reduce net production, and high *G*_*M*_ should tell muscle to increase uptake. Further, these functions must be set (e.g., by slow adaptation) so that, in the fasting steady state, production balances total utilization, *m*_*L*,0_ + *m*_*K*,0_ = *R*_0_ (*SI Appendix*, Eq. (42)), of which muscle consumes *R*_*M*,0_ ≈ 0.2 *R*_0_ [27].

Now consider a bout of moderate exercise, long enough to deplete muscle-stored glycogen (e.g., 60 min). Wholebody glucose uptake then increases roughly three-fold [31], and this demand is met primarily by increased hepatic production [32, 33]. The system settles into a new steady state, denoted by ex, and we denote differences from rest conditions by Δ_ex_. In particular, the exercise-induced increase in muscle glucose demand is Δ_ex_*R*_*M*_ = *R*_*M*,ex_ − *R*_*M*,0_ ≈ 2*R*_0_. Although *mechanistically* muscle uptake must depend on *G*_*M*_, a failure to meet demand would be a failure of control, and we therefore optimistically assume perfect tracking *m*_*M*_ (*G*_*M*_) = − (*R*_*M*,0_ + Δ_ex_*R*_*M*_).

Since total utilization increases in exercise, hepatic production *m*_*L*_ also increases (*SI Appendix*, Eq. (42)). Using the steady-state form of Eq. (6), the exercise-induced change in the liver–brain glucose difference obeys

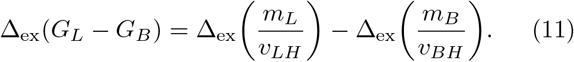

With Δ_ex_*m*_*L*_ *>* 0, Δ_ex_*m*_*B*_ ≈ 0, and fixed flow rates, the liver–brain gradient must increase. In reality, this effect is even larger: during exercise *v*_*LH*_ decreases substantially to support the increase in *v*_*HM*_, which provides oxygen to the working muscle, while brain flow increases modestly. Estimating a 20% increase in *v*_*BH*_ [34] and a 40% decrease in *v*_*LH*_ [35] yields an even larger Δ_ex_(*G*_*L*_ − *G*_*B*_).

Because *m*_*L*_(*G*_*L*_) must be a decreasing function, an increase in hepatic output necessarily implies a *decrease* in *G*_*L*_, so Δ_ex_*G*_*L*_ *<* 0. Thus, the increased liver–brain difference (Eq. (11)) produces a decrease in brain glucose,

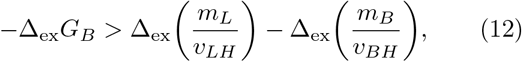

which can be large, especially with flow changes. Thus, in any hormoneless instantaneous local control scheme, increased muscle demand inevitably forces a drop in steadystate brain glucose. Figure 2(a) illustrates the minimal drop in *G*_*B*_, Eq. (12), during exercise for fixed flows (blue) and exercise-modified flows (red). The black dot indicates our reference demand Δ_ex_*R*_*M*_ ≈ 2*R*_0_.

**FIG. 2.**
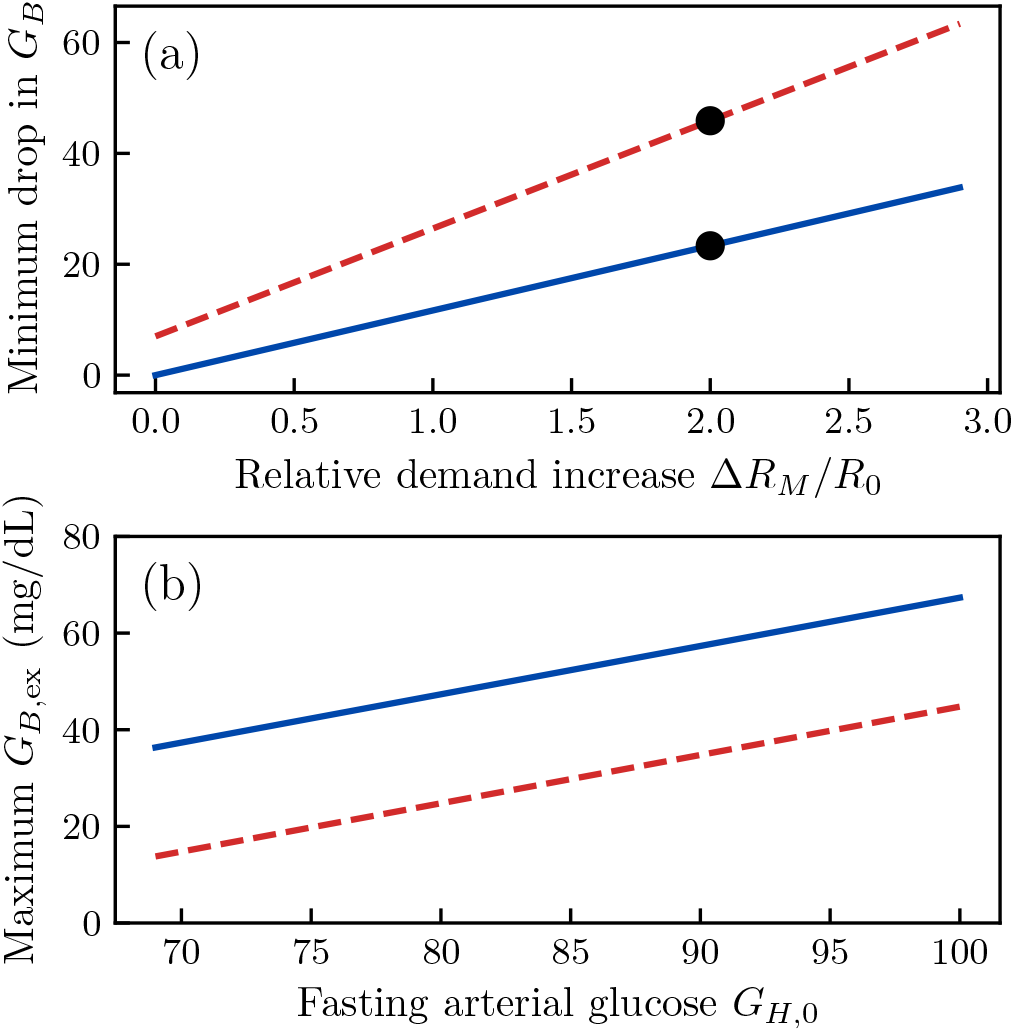
Exercise produces brain hypoglycemia in hormoneless control: (a) As the glucose demand imposed by muscle increases, brain glucose must drop. (b) Resulting trade-off between fasting arterial glucose and the highest achievable brain glucose during exercise. In both panels, solid blue curves use resting flow rates; dashed red curves use a 20% increase in brain blood flow and a 40% decrease in liver blood flow.

This drop in *G*_*B*_ implies a trade-off: either maintain normal fasting glucose and tolerate low brain glucose during exercise, or elevate fasting glucose to provide a safety margin for exercise. Figure 2(b) shows the Pareto front: the highest achievable *G*_*B*,ex_ as a function of fasting *G*_*H*,0_ for Δ_ex_*R*_*M*_ ≈ 2*R*_0_. Neurological symptoms begin near arterial glucose *G*_*H*_ ≈ 55 mg*/*dL [6], corresponding to *G*_*B*_ ≈ 46 mg*/*dL. We see that, with realistic blood flow changes, this threshold is only met in exercise for *G*_*H*,0_ *>* 98 mg*/*dL: right at the 100 mg*/*dL American Diabetes Association definition of impaired fasting glucose. A biochemically realistic curve *m*_*L*_(*G*_*L*_) would worsen this trade-off. Thus, with hormoneless instantaneous control, hypoglycemia during exercise cannot be avoided without chronically elevated fasting glucose.

Finally, recall that we assumed that muscle uptake always matches demand, regardless of local glucose. In reality, uptake must depend on *G*_*M*_ . Muscle glucose would also drop in exercise under hormoneless control, despite the increase in *v*_*HM*_ (*SI Appendix*, Section S5 B). Thus, hormoneless control not only risks brain hypoglycemia, but may also fail to meet demand in the muscle.

#### 2. Hormoneless control imposes a trade-off between hyperglycemia in a meal and flexible delivery of the meal to different organs

During the digestion of a meal, nutrients are released from the gut, blood glucose is elevated, and muscle and liver increase glucose uptake. Diabetes research focuses on how the body prevents hyperglycemia (high blood glucose) in meals, but glucose is *a fuel*, and the body should also be able to control each organ’s uptake to meet current needs.

We again study steady states, under a carbohydrate “ingestion rate” *m*_meal_, of which fraction 1 − *χ* is glucose. Thus, Δ_meal_*m*_*U*_ = (1 − *χ*)*m*_meal_. Other carbohydrates are converted to glucose in the liver, so the remaining *χm*_meal_ contributes to liver glucose flux (see below). Constant *m*_meal_ is a reasonable proxy for a slowly digested meal, and in healthy humans intravenous infusions of magnitudes *R*_0_– 2*R*_0_ reach steady states in around an hour [36], justifying this choice. Again, we expect non-constant *m*_meal_(*t*) to make control harder, only strengthening our results.

If a meal is eaten after exercise, more of the glucose load is taken up by muscle [37]. Since exercised muscle needs to replenish depleted glycogen, we postulate that this uptake fraction is *controlled*. Thus, in hormoneless control, we ask whether the muscle can control the value of *ϕ* ≡ − Δ_meal_*m*_*M*_ */m*_meal_, up to some value *ϕ*_max_.

In principle, the liver response may depend both on local glucose and on the local concentration of other carbohydrates; we thus model it as Δ_meal_*m*_*L*_(*G*_*L*_, *χm*_meal_).

The muscle can only attempt to control *ϕ* by adjusting *m*_*M*_ (*G*_*M*_). This control should be robust to meal composition and the rate of gastric emptying; we thus ask that it is robust to *χ* and *m*_meal_. We show in the *SI Appendix* (Section S5 C) that this requires

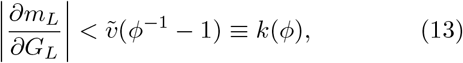

where 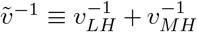. This is because making *ϕ* independent of *m*_meal_ requires the muscle to invert *G*_*M*_ (*m*_meal_). Since *G*_*L*_ − *G*_*M*_ ∝ *m*_*L*_, if 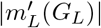 is too large *G*_*M*_ (*m*_meal_) is nonmonotonic, and this inference fails.

Thus, requiring that *ϕ* can be controlled limits the “strength” of the liver response to hyperglycemia, resulting in a lower bound on Δ_meal_*G*_*H*_ . The bound depends on the “operating range” over which muscle can control *ϕ*: the maximum *ϕ*_max_ that can be achieved, and the maximum *m*_meal,max_ for which it can be achieved. Defining M = (1 − *ϕ*_max_)*m*_meal,max_, we show (*SI Appendix*, Section S5 C)

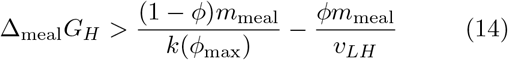

when (1 − *ϕ*)*m*_meal_ ≤ ℳ, and

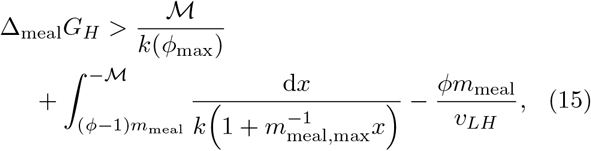

when (1 − *ϕ*)*m*_meal_ *>*ℳ.

To estimate a plausible *m*_meal_, note that a person needs *R*_0_ × 24 hr ≈ 200 g of glucose per day. This could come from stored carbohydrates from three meals, each slightly smaller than a standard 75 g oral glucose tolerance test (OGTT). In an OGTT, glucose leaves the gut over roughly 4 hours [38]. Thus, we split the daily demand over 12 hr, giving *m*_meal_ ≈ 2*R*_0_. In an OGTT, the peak glucose appearance rate is roughly double the average rate [38], so a reasonable *upper* limit is *m*_meal_ ≲ *m*_meal,max_ = 4*R*_0_. For a meal after an overnight fast, *ϕ* ≈ 20%–40% [39, 40]. Since an exercised muscle can take up ∼120% more glucose in a meal [37], we take *ϕ*_max_ = 0.75. Figure 3 then shows the minimum meal hyperglycemia for *m*_meal_ = 2*R*_0_ (blue) and 4*R*_0_ (red). The black dot is an estimate of the peak glucose in a healthy OGTT (a broad peak around 45–60 minutes) [38], placed at the estimate *ϕ* ≈ 0.3. We see that humans surpass the performance achievable by hormoneless control.

**FIG. 3.**
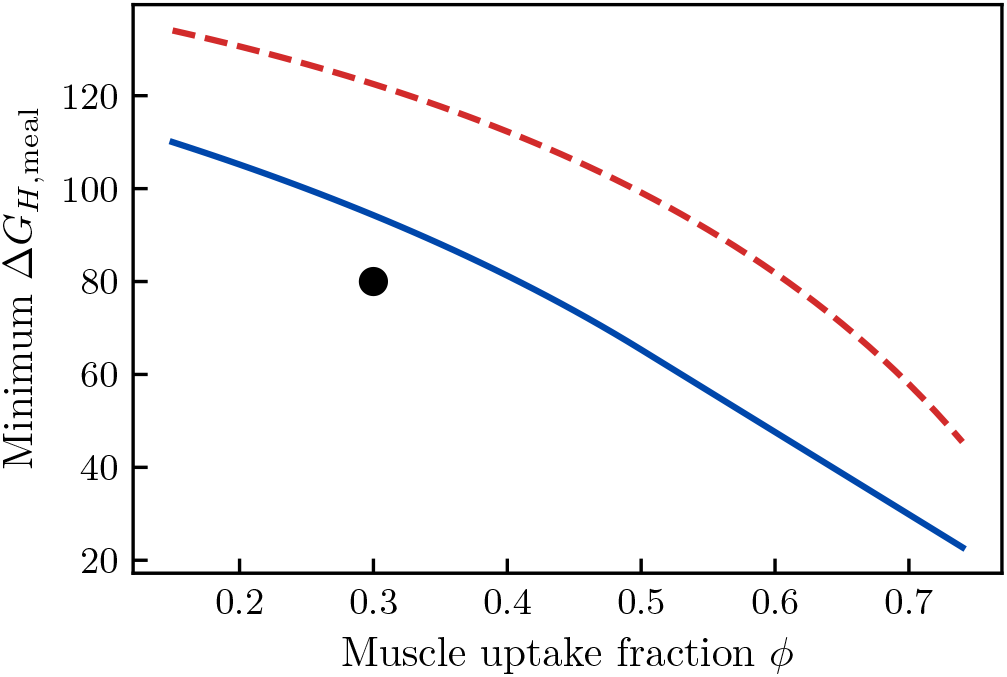
Minimum effect of a carbohydrate meal on arterial blood glucose under hormoneless control: Here, liver response is set to allow the periphery to “demand” any *ϕ* ≤ *ϕ*_max_ = 0.75 for any “meal rate” less than 4*R*_0_. The solid blue line shows the lowest achievable *G*_*H*,meal_ for *m*_meal_ = 2*R*_0_, in steady state, and the dashed red line shows the same for *m*_meal_ = 4*R*_0_ (conservative and aggressive estimates of the absorption rate in an OGTT). Both curves show greater hyperglycemia than a normal, healthy peak blood glucose in an OGTT (black dot).

#### 3. Hormoneless control is not robust to changes in blood flow

Blood flow rates change substantially on short timescales. For example, changes in posture [41] and circadian rhythms [42] can each produce ∼25% changes in liver blood flow. This produces changes in the liver– heart glucose difference that any control algorithm must deal with.

Consider the steady state produced by a change in *v*_*LH*_ during fasting. Since liver and muscle are most sensitive to changes in glucose, we assume that only *m*_*L*_ and *m*_*M*_ change. Thus, steady state is determined by the equations:

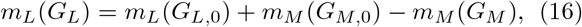

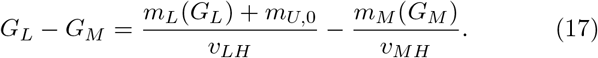

Suppose *v*_*LH*_ decreases. If *G*_*L*_ did not change, then Eq. (17) implies that *G*_*M*_ would decrease. This would decrease muscle glucose uptake, raising *G*_*L*_. Thus, a decrease in *v*_*LH*_ increases *G*_*L*_ and decreases *G*_*M*_, while an increase in *v*_*LH*_ does the opposite.

We write 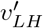 for the altered liver blood flow and denote changes produced by it with Δ_*v*_. It is undesirable for the muscle glucose uptake to fluctuate dramatically due to changes in liver blood flow. Thus, the body should care about the value of 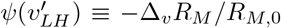. Defining an “effective stiffness” 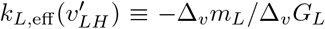, a short calculation shows

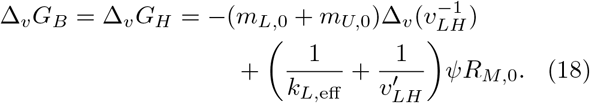

A particular design *m*_*L*_(*G*_*L*_) and *m*_*M*_ (*G*_*M*_) will result in particular functions 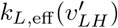 and 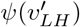.

The first term in Eq. (18) comes from the change in the total gradient, shown in Fig. 4(a). If *m*_*L*_(*G*_*L*_) turns off rapidly for *G*_*L*_ *> G*_*L*,0_, then *G*_*L*_ will rise very little, and thus the larger gradient will result in a drop in brain glucose. This suggests a trade-off between robustness to changes in *v*_*LH*_ and controlling hyperglycemia.

**FIG. 4.**
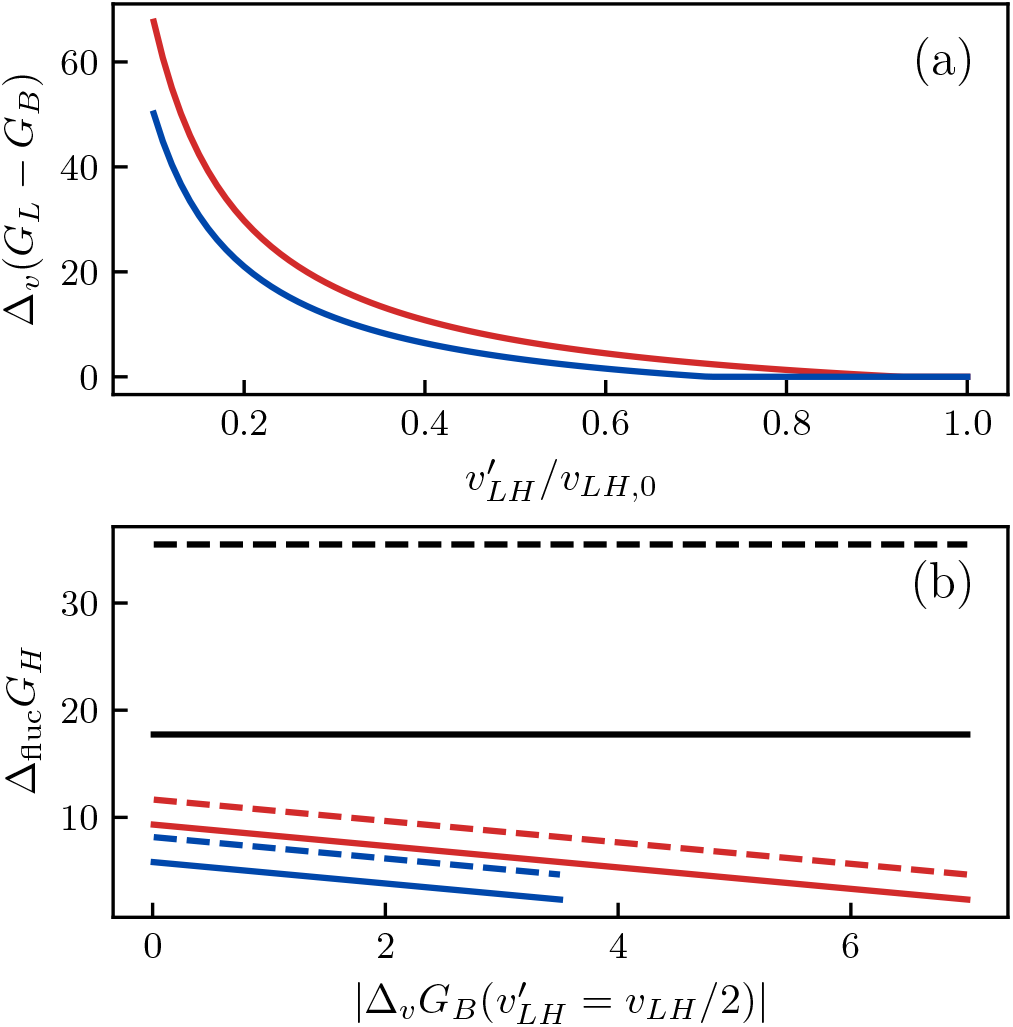
Effect of liver blood-flow change on fasting glucose in instantaneous, hormoneless control: (a) A decrease in liver blood flow increases the liver–heart glucose difference, whose magnitude is larger if the muscle response is tuned to avoid too large a fractional change *ψ* in glucose uptake (blue: *ψ* = 0.25, red: *ψ* = 1.0). (b) Trade-off between the minimum drop in brain glucose induced by a transient 50% drop in liver blood flow and the minimum increase Δ_dem_*G*_*H*_ produced by a 20% (solid) or 40% (dashed) drop in glucose demand; color indicates *ψ* as in (a). Black indicates the larger minimum Δ_dem_*G*_*H*_ imposed if the requirement that *ϕ*_max_ can be achieved (see discussion of meals) is imposed all the way down to meals smaller than Δ_dem_*R*.

In particular, note that glucose demand fluctuates over hours–days, necessitating changes in liver glucose output. Total glucose flux drops nearly a third overnight [43], and fasting glucose production can change by 40% with a 5-day change in diet composition [44]. In real humans, these changes may be driven partly by hormones, but they presumably reflect underlying changes in *demand*, which hormoneless control would also need to meet.

As we vary the total peripheral demand by some amount Δ_dem_*R*, the associated Δ_dem_*G*_*L*_ will also define a *k*_*L*,eff_ (Δ_dem_*R*) ≡ −Δ_dem_*R/*Δ_dem_*G*_*L*_. In particular,

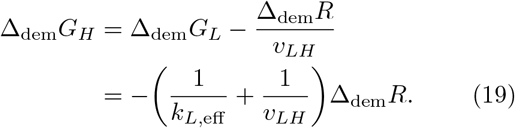

When Δ_dem_*R* = − *ψR*_*M*,0_, the two definitions of *k*_*L*,eff_ coincide, producing a trade-off between Δ_dem_*R* and |Δ_*v*_*G*_*B*_ | (calculations in *SI Appendix*, Section S6). Figure 4(b) shows this trade-off for a 50% drop in *v*_*LH*_ and Δ_dem_*G*_*H*_ for a 20% (solid) or 40% (dashed) fluctuation in demand.

Since Δ_*v*_*G*_*B*_ only constrains *k*_*L*,eff_ for |Δ_dem_*R*| = *ψR*_*M*,0_, the solid blue, dashed blue, and dashed red curves in Fig. 4(b) are loose bounds, which are only saturated if *m*_*L*_ suddenly decreases very rapidly above Δ_*v*_*G*_*L*_. We suspect a more realistic model of the dynamics would create a further trade-off, where such a rapid change causes instability; see *Discussion*, below.

If we require *ϕ* to truly be controllable for *any* meal, even those with *m*_meal_ ≪ *R*_0_, then the upper bound on *k*_*L*,eff_ required to achieve *ϕ*_max_ = 0.75 is smaller than that required to control Δ_*v*_*G*_*B*_, and thus Δ_dem_*G*_*H*_ would actually be determined by a trade-off with *ϕ*_max_, as shown by the black lines in Fig. 4(b).

### E. More complex control schemes using the history of local glucose still fail

So far we have assumed that each organ responds to its *current* local glucose. A natural generalization is for an organ to respond to the history of its local glucose, which it could do by accumulating an intracellular signal over time. In control-theoretic terms, the schemes considered above are nonlinear versions of proportional control, and the standard linear extension is proportional–integral– derivative (PID) control, which is also the basis of several commercial artificial pancreas systems [45]. Could such a scheme fix the problems of local control?

The failures of hormoneless control that we identified arise already in steady state, so we continue to analyze this limit (which we recall is an optimistic best case, since transient demands are harder to meet than sustained ones, and the transport delays we neglect would further degrade performance). In this limit, the derivative term in a PID controller has no effect since d*G*_*i*_*/*d*t* = 0. Thus, the question is whether PI control, or something similar, improves on control using current glucose alone.

Conventionally, by a PI controller we might mean

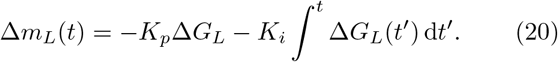

For many other problems, this controller is ideal, because it enforces Δ*G*_*L*_ = 0 at long times without producing the instabilities associated with high *K*_*p*_. However, it does not address the failures we have identified. Integral action drives Δ*G*_*L*_ → 0, so that −Δ_ex_*G*_*B*_ = −Δ_ex_(*G*_*L*_ − *G*_*B*_), and thus the exact same drop in brain glucose identified above still occurs. In a meal, a large *K*_*i*_ prevents the body from controlling *ϕ* just as a large *K*_*p*_ does. Conventional PID control therefore does not solve the problems of control using local glucose. We now consider a less conventional scheme, designed specifically to control *G*_*B*_ during exercise.

It seems that perhaps the liver could use the fact that it “knows” its own glucose production rate *m*_*L*_(*t*) to construct an estimator *Ĝ*_*H*_ of *G*_*H*_ from *G*_*L*_. It can then adjust *m*_*L*_(*t*) until *Ĝ*_*H*_ = *G*_*H*,0_, the fasting arterial concentration. Such a scheme would look like

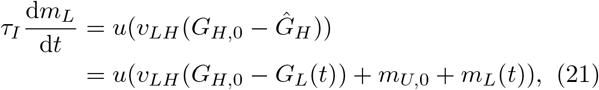

for some increasing function *u* with *u*(0) = 0.

Indeed, this scheme *does*, at first glance, seem to prevent brain hypoglycemia in exercise. Steady state requires d*m*_*L*_*/*d*t* = 0, giving *G*_*H*_ = *G*_*H*,0_ as long as *m*_*U*_ = *m*_*U*,0_. This more complex hormoneless control scheme, however, has unacceptable flaws. Firstly, *G*_*H*_ is still not robust to changes in *v*_*LH*_, unless the liver is assumed to “know” the blood flow rate 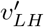 and change its behavior accordingly. Secondly, this scheme is not robust to variation in *m*_*U*,0_. Finally, the steady state in this scheme is unstable. For any value of *τ*_*I*_ and any *u*, a numerical calculation shows that such a scheme is linearly unstable, and introducing a simplification of the model also allows an analytical argument that shows the instability should remain for all possible physiological parameters (*SI Appendix*, Section S8).

Thus, schemes using the past history of glucose cannot eliminate the failings of local control in steady state. Integral control may still be necessary for a different reason. In a model resolving spatial structure within organs, stiff proportional responses may become unstable, and integral action is the standard remedy (see *Discussion*). Integral action also makes the effective liver stiffness *k*_*L*,eff_ depend on timescale, so a liver that is stiff to slow fluctuations in demand and soft to rapid changes in blood flow could possibly evade the trade-off shown in Fig. 4(b). Quantifying this requires a frequency-resolved analysis, which we leave to future work.

### F. Hormonal control eliminates these trade-offs

We have shown that hormoneless control algorithms face severe tradeoffs. We now analyze hormonal control algorithms, and show that they can surpass the limits we derived for hormoneless control. We assume insulin is produced in the pancreas at rate *βf* (*G*_*P*_), is cleared in the liver at rate *γ*_*L*_*I*_*L*_, and directly controls liver glucose production *m*_*L*_(*I*_*L*_) [46, 47]. (Our conclusions still hold if insulin is cleared in multiple organs, but explicit formulas are no longer available.)

The insulin secretion rate is a function of *G*_*P*_ (*t*). In our theory, a crucial benefit to placing hormone-secreting cells in the pancreas is that the total glucose consumption rate of the pancreas is very small relative to blood flow. As a result, the equation for d*G*_*P*_ */*d*t* gives

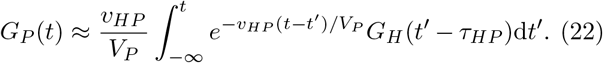

*τ*_*HP*_ and *V*_*P*_ */v*_*HP*_ are both only a few seconds. Thus, the insulin secretion rate *βf* (*G*_*P*_) in the pancreas responds to the past few seconds of *G*_*H*_ (*t*), with a few-second delay.

Insulin in target tissues thus reflects the history of *G*_*H*_ (*t*) (details in *SI Appendix*, Section S7). Thus, hormonal control allows all organs to respond to the history of *G*_*H*_ (*t*), rather than the history of their local glucose concentrations. Responding to a centralized glucose rather than a local one reduces the effect of glucose gradients, and this eliminates the trade-offs that we have identified. For example, the steady state in exercise obeys (*SI Appendix*, Section S7)

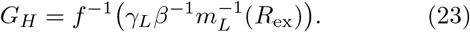

By choosing *m*_*L*_(*I*_*L*_) and *f* (*G*_*P*_) to be strongly varying, and thus making their inverses vary slowly, evolution can make the dependence of *G*_*H*_ on *R*_ex_ arbitrarily weak. Thus, a hormone can eliminate the trade-off between fasting hyperglycemia and exercise hypoglycemia.

Similarly, at steady state during a meal, we can determine what is achievable using hormones by allowing both liver and muscle to respond to *G*_*H*_, rather than *G*_*L*_ and *G*_*M*_ respectively. Since the muscle and liver respond to the same signal, it is straightforward for the muscle to control *ϕ* as long as it knows how the liver response function is programmed (*SI Appendix*, Section S7). Thus, the trade-off shown in Fig. 3 is completely eliminated.

Although our estimate for the typical peak hyperglycemia in an OGTT is lower than would be achievable with hormoneless control, it is still of a similar magnitude, despite the fact that hormones could completely eliminate the trade-off. Humans therefore do not push meal hyperglycemia as low as hormones allow. Mechanistically, this probably reflects the fact that liver and muscle responses in meals depend on local glucose, not just hormonal signals [48]. From an adaptive perspective, it may hint at another unknown control objective that is simultaneously being optimized. It may also reflect some limitation of our approximations (e.g., working in steady state).

Finally, we note that the steady-state equations for *G*_*H*_ with hormones (*SI Appendix*, Section S7) have no dependence on *v*_*LH*_, so the sensitivity to blood flow changes is also completely eliminated.

## III. DISCUSSION

In a simplified model of physiological fluxes and blood flow, we have shown that control schemes where each organ responds directly to local glucose face unavoidable trade-offs: between fasting normoglycemia and preventing brain hypoglycemia in exercise, between hyperglycemia and flexible allocation of glucose in meals, and between robustness to blood flow and demand changes. We have further shown that a coordinating signal, such as a circulating hormone, can eliminate these trade-offs.

Existing experimental data are consistent with these results [32, 33]. The main triggers of increased glucose production in exercise are insulin decrease, glucagon, and catecholamines. Human exercise experiments that clamp insulin and glucagon, while blocking adrenergic action, so that tissues are mostly left to regulate based on the glucose they see, observe a ∼52 mg*/*dL fall in blood glucose [31, 49]. Thus, the body clearly relies on hormones to regulate glucose production in exercise, as our theory says is necessary. The numerical value is also consistent with our *>* 44 mg*/*dL expectation for hormoneless control strategies. This is, however, not a full confirmation of our analysis since the organ responses of these subjects are tuned for a body that has insulin, and were never adapted to work without it.

Notice that our analysis of the performance achievable using insulin in exercise also applies to glucagon, since both are secreted in the pancreas.

In contrast, glucose uptake in hyperglycemia responds to *both* glucose and insulin [48]. As noted above, this is likely connected to the fact that human physiology does not push meal hyperglycemia that much lower than would be achievable without hormones, and may reflect the existence of another, not-yet-identified control objective that is being optimized, which would impose new tradeoffs on the objectives that we considered here.

### A. Evolutionary and physiological consequences

Our analysis suggests possible functional advantages for a variety of other features of human anatomy and physiology, as well as for the physiology of non-human animals.

Firstly, our analysis explains where in the circulation insulin must be secreted, since the beta cell has to sample blood whose glucose reflects *G*_*H*_ . If the islets were located *after* the liver, then the glucose they sense, and thus *I*_*L*_, would reflect *G*_*L*_ rather than *G*_*H*_, and the drop in blood glucose in exercise would remain. If they were instead downstream of the gut, a meal being slowly digested would interfere with the response to exercise. The same argument should apply to glucagon, which is secreted from the same organ. More speculatively, requirements of this kind may help explain why the blood circulation in humans is organized in the topology it has, with the gut and the pancreas in parallel with each other, and in series with the liver, instead of on dedicated branches.

Next, we note that long-range signaling is not the only conceivable way out of these trade-offs, which arise from the finiteness of blood flow rates. Evolution could have relieved them by increasing blood flow instead. That it did not implies further trade-offs between glucose regulation and whichever costs drove the evolution of the cardiovascular system, which presumably also constrain how blood flow is allocated during exercise.

A related evolutionary concern is that, although details vary substantially, insulin plays a role in glucose regulation across the vertebrates [50, 51]. It would therefore be concerning if the benefits we ascribe to hormonal glucose control only applied in humans. Crucially, the allometric scalings of cardiac output and metabolic rate are similar so that the oxygen demand can be met with only the necessary cardiac effort [52]. We would expect the glucose demand to broadly scale in proportion to the metabolic rate, and flow rates to each organ to scale in proportion to cardiac output. This suggests that trade-offs similar to those we have identified are relevant in vertebrates at all scales, consistent with such trade-offs being among the crucial adaptive benefits of hormonal glucose control.

Another consequence of our results is for the need to maintain brain blood flow in exercise. Here, muscle blood flow increases to meet oxygen demand, and blood flow to the gut and liver decreases to help accommodate this increase. Since brain blood flow is more than enough to meet its oxygen demand, it is surprising [53] that this flow does not also decrease to help supply oxygen to muscle. Our results reveal a key reason: when glucose is controlled by pancreatic hormones, *G*_*B*_ is independent of *v*_*LH*_, but remains sensitive to *v*_*BH*_ .

Our analysis of meals was built on a putative trade-off between hyperglycemia and control of the fraction *ϕ* of the meal which is taken up by the muscle (or more broadly, the periphery). Whether the body can control *ϕ* robustly under different conditions is unknown, but our theory predicts that, without the need for such control, the body would be able to control hyperglycemia much better. This thus opens possibilities for experiments to directly test if the body tries to robustly control the fraction *ϕ* of a meal taken up by each tissue.

Finally, we remark that the intimate connection between glucose control and cardiovascular physiology in our theory may have a variety of important consequences. Glucose gradients also become larger when the circulation itself is compromised, and this likely constrains human physiology. For example, in critical illness, impaired cardiac function would increase glucose gradients, so holding arterial glucose fixed could risk brain hypoglycemia. This may be an adaptive reason for stress hyperglycemia [54]. Notice also that cardiovascular symptoms are some of the main complications of diabetes. It is thus tempting to speculate that there may be a positive feedback loop between hyperglycemia and cardiovascular damage.

### B. Limitations and future work

In principle, a pancreatic hormone is not the *only* mechanism which can coordinate organs to respond to *G*_*H*_, eliminating the trade-offs we have identified. Neural signaling could achieve the same coordination. More speculatively, since each organ receives some arterial blood input, one could imagine a design where a specialized cluster of cells on the arterial end of each organ sensed glucose, and then released a paracrine or shorter-range endocrine signal that traveled through the organ and produced an appropriate response. Such systems presumably would have their own advantages and disadvantages relative to a single central hormonal signal, which would be interesting to study in future work.

Since time delays can produce instabilities with high-gain control [55], one might have expected such instabilities in our models. However, we find that this effect is not present in our model, because we have assumed that each organ is well-mixed (*SI Appendix*, Section S9). In a model with spatial structure within organs, we expect that instability with high-gain control would impose further limits on 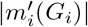, further worsening the performance of local control (although this problem might be solved by strategies like integral control, as mentioned above).

Our whole-body glucose allocation control framework may be used to analyze other physiological states. In the current analysis, slow changes in glucose demand, e.g., the 20–25% increase over the course of pregnancy [56], could be compensated by local adaptations. Further, increases in the glucose gradients between the liver, heart, and fetus could already be prevented by the observed dramatic increases in blood flows to the liver [57] and the uterus [58]. Thus, understanding the adaptive benefit of control of insulin sensitivity by placental lactogen hormone is a question for future work.

Relatedly, many other long-range signals modulate effects of insulin on glucose dynamics. Examples include glucagon [59], incretin hormones [60], and glucose sensing in the portal vein [61]. These provide additional information, often anticipatory, about local glucose concentration or the expected demands in different organs, and would need to be analyzed with more extensive, spatiotemporally complex control objectives in mind.

### C. Potential clinical impact

In the modern world, diabetes is an enormous problem. It is thus tempting to view avoiding hyperglycemia as the sole goal of glucose control. We have shown that other goals—meeting each organ’s glucose needs, robustness to blood flow changes, and preventing demand in one organ from producing low glucose in another—pose greater challenges, and trade off against preventing hyperglycemia. While, in our analysis, we focused on counterfactuals, understanding why the body is designed the way it is may have a substantial biomedical impact. Indeed, what a control system *is for* determines *how it breaks*, so our analysis should help explain the pathophysiology of diabetes and guide its treatment.

Improving artificial glucose control systems may be one of the first applications of these ideas. While closedloop systems are now common in clinical applications, they still perform worse than a healthy pancreas [45], and exercise is among the hardest conditions for them to respond to [62, 63]. Some of that difficulty may come from such artificial systems having to solve the same tradeoffs that we analyze here, and, in particular, from where they measure glucose. A subcutaneous sensor sits in a peripheral tissue with its own arteriovenous difference, so it does not report the arterial concentration *G*_*H*_ that the pancreas responds to (this is in addition to the well-known delay between subcutaneous and plasma glucose). At rest, the gap is a few mg/dL, possibly smaller than the sensor error. It may widen during exercise, when liver and muscle fluxes increase dramatically: this change is likely unpredictable, due to its sensitivity to muscle blood flow changes. Aortic sensing is not an option, so the practical question is whether the gap can be estimated instead. Several sensors at different sites, read together with a measure of activity, might be enough to reconstruct *G*_*H*_ . Designing such practical algorithms is a promising direction for future work.

## ACKNOWLEDGMENTS

We thank Helen S. Ansell, K. M. Venkat Narayan, Alejandro Caicedo, Darko Stefanovski, and Cora Hirst for helpful discussions and/or reading drafts of this manuscript. This research was supported in part by grants from the NSF (DMS-2235451) and Simons Foundation (MPS-NITMB-00005320) to the NSF-Simons National Institute for Theory and Mathematics in Biology (NITMB). IN and SAR were supported, in part, by the Simons Foundation Investigator Program grant to IN. SAR was additionally supported by Emory Global Diabetes Research Center and Emory University Research Council grants to IN and PV. PV was supported, in part, by the NIH through grants 5P30DK125013 and R03DK129627. Part of this work was performed at the Aspen Center for Physics, which is supported by National Science Foundation grant PHY-2210452, and IN thanks the Center for the time and quiet to think. Our numerical calculations for this work (including larger-scale exploratory simulations) were managed using signac [64]. We acknowledge support of our work through the use of the HyPER C3 cluster of Emory University’s AI.Humanity Initiative.

## SUPPORTING INFORMATION

### S1. THE MINIMAL MODEL OF GLUCOSE HOMEOSTASIS ALLOWS EQUIVALENT HORMONELESS CONTROL

The system for *I*(*t*) and *I*_2_(*t*) in the main-text model, Eqs. (1)–(3), can be formally solved by treating *G*(*t*) as a given function and applying variation of parameters. This gives for *I*(*t*):

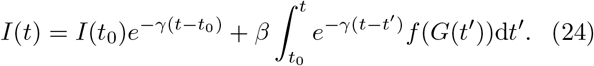

The exact same method applied to the equation for *I*_2_(*t*) gives:

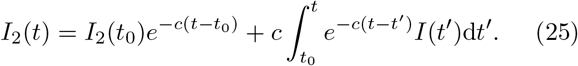

As long as *t* ≫ *t*_0_, so that the system forgets its initial conditions, the first term in each of these equations, representing transient history of the initial conditions, becomes negligible. Imagining that the system has been running forever then yields the compact formula for *I*_2_(*t*) given in the main text, Eq. (4).

### S2. OTHER WELL-MIXED AND COMPARTMENTAL MODELS ALSO ALLOW FOR HORMONELESS CONTROL

Here we briefly comment on other types of models that can be rewritten as hormoneless control algorithms.

#### A. Calcium homeostasis

A classic paper [1] argues that parathyroid hormone and 1,25-dihydroxycholecalciferol implement integral feedback of plasma calcium. Because their model is equivalent to a PI controller acting on plasma calcium, it is a special case of a model that responds to the history of that concentration.

#### B. A simple insulin–glucagon model

A recent paper argues for functional benefits of the paracrine interactions between insulin *I* and glucagon *C* [2] using the model:

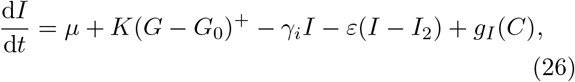

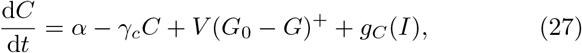

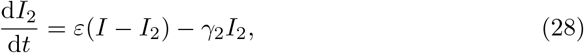

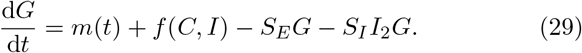

It focuses on the case where *g*_*I*_ and *g*_*C*_ are linear. In this case, the explicit mapping of the main text is straightforward to generalize. The variable **z** = (*I, I*_2_, *C*) satisfies a linear inhomogeneous ODE

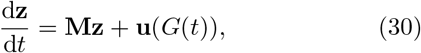

and therefore, under the assumption that **M** has only negative eigenvalues (necessary for stability), it admits an explicit solution

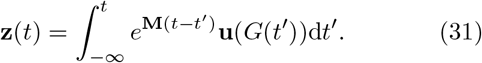

Thus, just as in the main text, the dynamics of *G*(*t*) can be rewritten as an explicit functional of the history *G*(*t*^*′*^ ≤ *t*).

Our logic also applies to the general case, without an explicit construction. Although *G*(*t*) is itself set by the coupled dynamics, we may regard it as a prescribed input and ask how the hormones respond. The state **z**(*t*) then obeys ODEs of the form

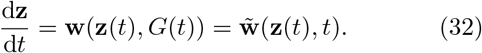

Under restrictions on *g*_*I*_ and *g*_*C*_ to prevent solutions from blowing up in finite time, the Picard–Lindelöf theorem [3] guarantees the existence and uniqueness of a solution **z**(*t*) to this ODE, after specifying *G*(*t*^*′*^ ≤ *t*), for a particular **z**(0). Assuming that the dynamics has a limited long-time memory, this precisely means that *I*(*t*), *I*_2_(*t*), and *C*(*t*) can be reconstructed deterministically from the trajectory *G*(*t*). For any particular *g*_*I*_ and *g*_*C*_, there exists a causal functional that allows the dynamics of *G*(*t*) to be written in terms of the past history *G*(*t*^*′*^ ≤ *t*).

#### C. More general well-mixed models of hormone networks

The final argument of the previous section is generic enough to apply to any model where a network of well-mixed hormones interacts with each other and a well-mixed regulated variable.

#### D. More complex compartmental models of glucose homeostasis

The UVA/Padova model [4] is a detailed compartmental model of glucose homeostasis in type 1 diabetes, accurate enough that the FDA has accepted it in place of animal trials for preclinical testing of glucose-control algorithms [5].

This model, as a model of T1D, lacks insulin secretion. Nonetheless, we may ask if, when supplemented with an insulin secretion model similar to those we have discussed, it would still describe a physiological control algorithm that is equivalent to a hormoneless algorithm. We find that control in this model is not strictly equivalent to any hormoneless algorithm, but that this model still cannot adequately describe the distributed control problems that we identify in the main text.

Specifically, glucagon *C* in this model has simple, single-compartment kinetics, with secretion rate that is a function of plasma glucose *G* and its time derivative. The glucose kinetics of the model contain two compartments, plasma and tissue

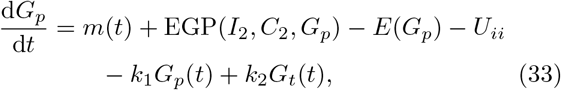

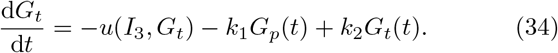

Here EGP is endogenous glucose production in the liver, controlled by the hepatic insulin- and glucagon-action variables *I*_2_ and *C*_2_. The rate of insulin-dependent glucose clearance *u* (e.g., in muscle) is controlled by the peripheral insulin-action variable *I*_3_, and *E* represents kidney glucose clearance.

*I*_2_ and *I*_3_ have simple exchange kinetics with the plasma insulin as in the main text. *C*_2_ has slightly more complicated kinetics, which only responds to elevation of *C* above basal (representing an effective response rather than simply diffusion into another compartment). Thus, as in the main text, *I*_2_, *C*_2_, and *I*_3_ simply reflect the history of plasma glucose (and exogenous insulin), if the model is supplemented with a standard insulin secretion model. As a result, glucose *production* in this model clearly reflects the history of plasma glucose.

Glucose *uptake*, however, cannot be rewritten this way. As Eq. (34) shows, uptake *u*(*I*_3_, *G*_*t*_) depends on the tissue glucose *G*_*t*_, a separate compartment from plasma glucose *G*_*p*_. Because *G*_*t*_ differs from *G*_*p*_ and is not determined by the plasma-glucose history alone, uptake cannot be expressed as a functional of that history. Yet, because production still reduces to the plasma-glucose history, the model lacks the spatial structure on the production (liver) side that drives the effects we identify in the main text, so it cannot reproduce them.

### S3. DETAILS OF FLOW MODEL

#### A. Time-delay approximation

Our results are obtained in steady state and do not depend on the dynamical details of the model. Nonetheless, we will briefly comment on our assumption that all species **x** are conveyed from organ *i* to organ *j* with a single time delay *τ*_*ij*_, to help guide future work analyzing dynamics in detail.

In reality, there is a distribution *P* (*τ*_*ij*_). Blood flow in the body spans from Re ∼ 10^−3^ in capillaries to nearly 10^4^ at peak systole in the aorta. These flows are not turbulent, so solute spreading can arise from two factors: (1) a broad velocity profile in the vessel (with solute at the edges traveling slower) and (2) lateral diffusion. Perpendicular diffusion blunts the first effect, by allowing solute in slow streamlines to move into fast ones [6]. Peclet numbers for glucose and insulin in large vessels are large, so lateral diffusion is likely to be unimportant.

The aorta sits at the high-Reynolds end of this range. Due to a combination of moderately high Reynolds number, pulsatile flow (large Womersley number, *α* ≈ 13– 21), and vessels shorter than the hydrodynamic entrance length, its velocity profile is blunt, with a boundary layer under 2 mm [7, 8]. Because the velocity profile is quite flat, the travel-time distribution is narrow and our approximation of a constant travel time is reasonable.

For laminar flow in small vessels, on the other hand, the distribution of travel times is actually quite broad at large Peclet number. Let *ū* be the mean flow velocity. At time *t*, the lateral spread of a drop of solute is 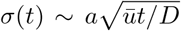 [6], and as *D* → 0 the variance of *τ*_*ij*_ diverges (in reality, the blood cells are expected to disrupt laminar flow, and branch points should also introduce some amount of turbulent mixing). Nonetheless, for simplicity we use a single travel time. The maximum rate of flow (at the center of a vessel) is 2*ū*. The mean travel time and the volumetric flow rate obey *v* = *πa*^2^*ū*, ⟨*τ* ⟩ = *L/ū*.

We suspect a broad distribution of travel times, in an analysis of short-timescale dynamics, would limit the speed of control: a wide distribution of travel times limits the reliability of responding aggressively to small, fast changes in glucose.

Because the most essential glucose dynamics are slower, minute–hour timescale ones, we expect the “well-mixed organ” approximation will be a more important flow-model approximation to improve in future work. In particular, in reality, much of the delay takes place “inside” the organ (in capillaries), rather than strictly “between” organs.

#### B. Model parameters

We have used a set of cardiovascular parameters that are representative of a resting adult, collected in Tables S1 and S2. We require the volumetric flow rates to satisfy volume conservation in each organ,

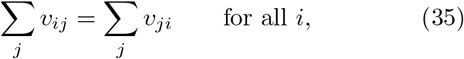

which is indeed consistent with the experimental data.

Total flow rates through an organ are unambiguous and model-independent, and can be confidently inferred from experiments, although reported values vary by ∼50% across studies, likely reflecting true physiological variation from person to person and condition to condition. The meanings of *V*_*i*_ and ⟨*τ*_*ij*_⟩, on the other hand, are somewhat model-dependent and thus cannot be uniquely related to experimental data. Generally, experiments measure the mean transit time of a tracer injected in one blood vessel, through an organ, to another vessel. Although much of this transit time is spent “in” the organ (certainly, in capillaries), we will attribute this entire delay to connections *τ*_*ij*_ in our model. When experiments only report mean transit times for multiple connections (e.g., heart → liver → heart), we will use measured mean blood velocities (and an assumption of equal length) to estimate the ratio of the components. For organ blood volumes *V*_*i*_, we take reference fractions assembled in [9] from many experimental sources, and multiply them by a reference blood volume of 5 L.

It is clear from the above discussion that we can be fairly confident in our assignment of typical flow rates *v*_*ij*_, and less confident in our determination of volumes *V*_*i*_, delays *τ*_*ij*_, and indeed the entire assumption that flow can be adequately modeled with a single delay. However, the results in the main text are obtained in quasi-steady states, which are determined solely by *v*_*ij*_. The detailed dynamics are sensitive to the values of *V*_*i*_ and *τ*_*ij*_, but these do not affect our results.

##### Flows to the liver

The liver is fed by the hepatic artery (directly from the heart; ∼250 mL/min [10]) and the portal vein (combining flows from gut, spleen, pancreas, and other sources; ∼950 mL/min [10]). Since the spleen accounts for a minuscule fraction of the body’s glucose utilization, however, we can treat flow through the spleen as a flow directly from the heart to the liver for our steady-state analyses. Thus, after subtracting estimated blood flows to the gut (∼400 mL/min [11]) and pancreas (∼150 mL/min [12]) (which both drain completely into the portal vein), we treat the remainder of the portal vein flow as a direct flow from heart to liver. This results in the flow rates *v* quoted in Table S1. To assign delays, we note that experiments have measured a mean hepatic artery → hepatic vein (“heart → liver → heart”) transit time of 26 seconds and a mean superior mesenteric artery (SMA) → hepatic vein (“heart → gut/pancreas → liver → heart”) transit time of 40 seconds [13]. Measured flow velocities in the hepatic artery and the hepatic vein are similar [14]; we thus split this delay evenly between the two. The pancreas and gut are fed and drained by the same main artery and vein, and we were unable to find (invasive) tracer measurements which would precisely measure the transit times through the pancreas in humans, so we assign the same delays for both routes, and split the delay evenly between the SMA and the portal vein. (Note that blood which passes through the spleen has a larger delay [13]; in future work requiring modeling dynamics, it may be necessary to add the spleen to the model.)

**TABLE S1.** Vessel parameters. Flow rates are derived from [10–12, 15, 18, 22, 23] and delays from [13, 14, 16, 17, 19– 21, 24, 25], as described in the text.

| Connection | Flow rate $v_{ij}$ | Delay $\tau_{ij}$ |
| --- | --- | --- |
| Heart → liver (including spleen) | 650 mL/min | 13 s |
| Liver → heart | 1200 mL/min | 13 s |
| Heart → pancreas | 150 mL/min | 13 s |
| Pancreas → liver | 150 mL/min | 13 s |
| Heart → gut | 400 mL/min | 13 s |
| Gut → liver | 400 mL/min | 13 s |
| Heart → brain | 750 mL/min | 3 s |
| Brain → heart | 750 mL/min | 6.5 s |
| Heart → muscle | 1000 mL/min | 20 s |
| Muscle → heart | 1000 mL/min | 40 s |
| Heart → kidneys | 1000 mL/min | 2 s |
| Kidneys → heart | 1000 mL/min | 2 s |

##### Flow to the brain

Using reported brain perfusion rates [15] and a rough brain mass of 1.4 kg gives an estimate of 750 mL/min. Cerebral artery flow velocities are around 40 cm/s, while jugular vein flow velocities are around 20 cm/s [16], so we estimate a mean transit time of a half-second from the heart to the internal carotid artery and one second from the internal jugular back to the heart. Flow from the internal carotid artery and internal jugular vein takes roughly 7.5 seconds [17], which we split between “heart → brain” and “brain → heart” using the same ratio as the flow velocities in the carotid artery and jugular vein, giving the values in Table S1.

##### Flow to the kidneys

Reported flow rates are roughly 1000 mL/min (combining both kidneys) in young and middle-aged subjects, although lower in elderly subjects [18]. The total renal circulation time is roughly 4 seconds [19]. Mean blood velocities in the renal arteries and veins are comparable [20, 21], so as a rough estimate we split this transit time evenly between heart→kidney and kidney→heart.

##### Flow to muscles

To obtain estimates for total blood flow to skeletal muscle, which is not all fed by a single artery or drained by a single vein, requires adding up estimates from various individual muscles. This results in estimates in the range of 750–1250 mL/min at rest [22, 23]; we take a value of 1000 mL/min. The time delay depends substantially on the specific element of muscle; for example, blood flow to the bottom of the leg involves a substantially larger delay than to the top of the leg; the mean of the two at low flow rate (rest) is roughly 20 s [24]. Venous flow returning from muscle is slower because the veins have larger diameters than the arteries. The femoral vein has roughly 150%–200% the cross-sectional area of the femoral artery [25], so we estimate the delay returning from muscle to be twice as large as the delay to muscle.

**TABLE S2.**
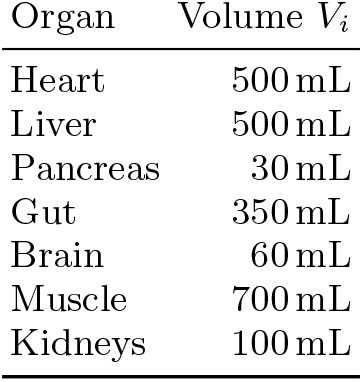
Estimated organ blood volumes. Reference fractions are taken from [9] and multiplied by a reference blood volume of 5 L, as described in the text.

### S4. MASS CONSERVATION

Here we briefly derive equations for the total amount **X**(*t*) of all species in the body within the model, and for its associated conservation law, which can simplify some derivations of steady-state equations in our model.

To compute the total amount of each species in the body in the model, we need to compute the amount contained within the blood vessels. Let **x**_*ij*_(*t, ℓ*) denote the concentration along the vessel from *i* to *j*, with *ℓ* ∈ [0, *L*_*ij*_] the position along the vessel. Let *A*_*ij*_(*ℓ*) be the cross-sectional area of the vessel. We then have

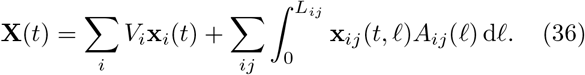

In our simple flow model, the flow speed *u*_*ij*_(*ℓ*) obeys *u*_*ij*_(*ℓ*) = *v*_*ij*_*/A*_*ij*_(*ℓ*). Thus,

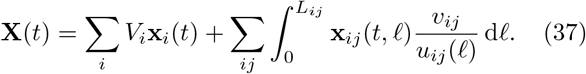

Now, define 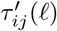 as the delay to reach position *ℓ* along the vessel; 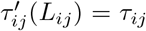. By this definition, 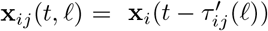. We further see that 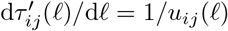, and thus

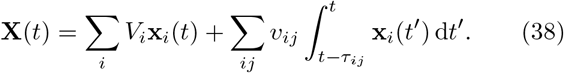

With this result in hand, we can obtain the conservation law for **X**. We have

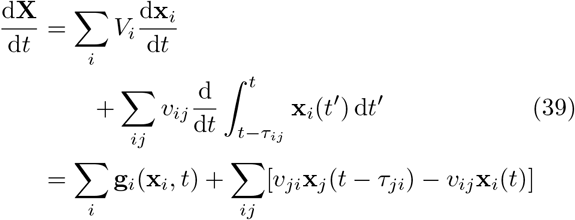

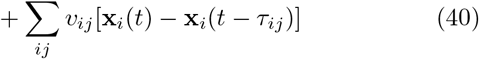

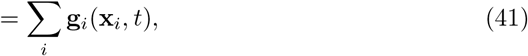

where the last line follows from breaking apart the first sum into two and relabeling dummy indices.

In steady state all concentrations are constant, so d**X***/*d*t* = 0 and

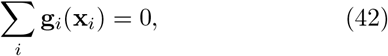

that is, production summed over all organs balances consumption. The transport terms cancel exactly despite the delays, which is what lets the calculations below impose total balance without tracking the glucose in transit through the vessels.

Equation (42) only holds if we include all organs—for glucose, this includes the blood/skin, adipose, and renal consumption/production mentioned in the main text, but often omitted in our calculations. However, notice that *changes* in **g**_*i*_ under some change in conditions must also sum to zero, and this is the fact that we use in practice in our derivations, considering a condition where only certain **g**_*i*_ change.

### S5. DETAILED STEADY-STATE CALCULATIONS

Here we provide full derivations of main-text results from Eq. (7) to Eq. (19).

#### A. Resting gradients

In steady state, the dynamics of the model, Eq. (6), reduce to

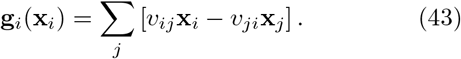

Presently, we only require the equations for the glucose component (as in the main text, we use *m* to denote this component of **g**). Organs with simple loop connections to the heart (muscle *M*, brain *B*, and kidneys *K*; see Fig. 1) yield very simple steady-state equations. Using volume conservation (Eq. (35)),

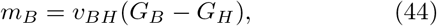

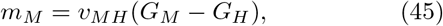

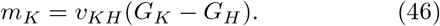

The liver gradient requires slightly more effort to obtain. Assuming glucose consumption in the pancreas is negligible, the steady-state equations for the gut, liver, and pancreas are:

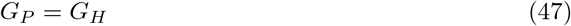

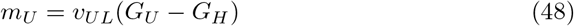

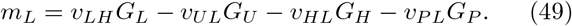

The first two equations can be used to eliminate *G*_*U*_ and *G*_*P*_ from the third, yielding

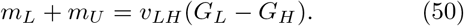

Thus, we have derived main text Eqs. (7)–(9).

#### B. Drop in muscle glucose in exercise under hormoneless control

Here, we present evidence for the main-text remark that “muscle glucose would also drop in exercise under hormoneless control, despite the increase in *v*_*HM*_ .” To see this, note that the *original* muscle–heart gradient at rest was quite small:

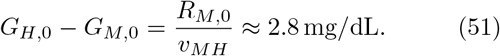

Muscle glucose uptake increases in exercise, which would tend to increase this gradient. Muscle blood flow increases as well, which would tend to decrease it. No matter how much blood flow increases, it cannot change the *sign* of this gradient:

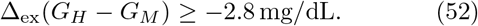

The drop in the heart is much larger than this for our reference exercise rate, even ignoring the change in liver blood flow:

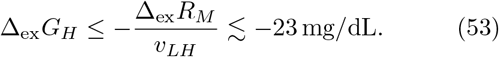

Thus,

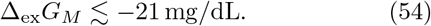

Since muscle uptake must in fact depend on *G*_*M*_, a drop of this size means our optimistic “perfect tracking” assumption is likely to fail, and hormoneless control would probably fail to meet glucose demand in muscle in exercise.

#### C. The trade-off between meal flexibility and hyperglycemia

As in the main text, we consider steady state for a “meal rate” *m*_meal_, of which a fraction 1 − *χ* is glucose and a fraction *χ* is other simple carbohydrates that are converted to glucose in the liver during the meal. Glucose absorption in the gut is thus

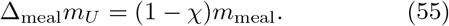

Glucose production in the liver is some combination of completely endogenous glucose production and conversion of simple carbs to glucose. The local concentration of carbohydrates can presumably be sensed, and the liver also “knows” how fast it is converting simple carbs to glucose. We thus allow a general control strategy

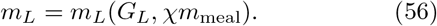

Our primary control objective is the ability of the muscle to *control ϕ*, in a way that is robust to *m*_meal_ or *χ*. Thus, muscle will choose some target *ϕ*_targ_, and choose *R*_*M*_ (*G*_*M*_, *ϕ*_targ_) such that *ϕ* = *ϕ*_targ_. How can it do so? Intuitively, if we assume that the muscle “knows” the dynamics of the rest of the body (i.e., the equations of our model), it can try to infer *m*_meal_ from local glucose measurements. We will find that achieving this requires two limits on the form of *m*_*L*_: it must depend additively on *χm*_meal_, and the magnitude of ∂*m*_*L*_*/*∂*G*_*L*_ cannot be too large. Thus, we will obtain a lower bound on Δ_meal_*G*_*H*_ .

We begin with the steady state equations for *G*_*M*_ and *G*_*L*_, as well as mass conservation, Eq. (42). These are:

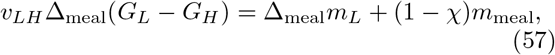

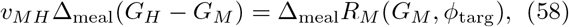

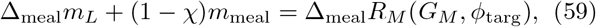

where *R*_*M*_ (*G*_*M*_) = −*m*_*M*_ (*G*_*M*_).

With a bit of algebra, these equations imply

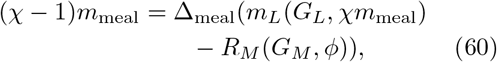

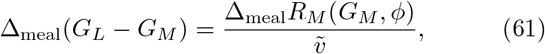

where we have defined 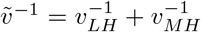. Now, substituting the second equation into the first yields

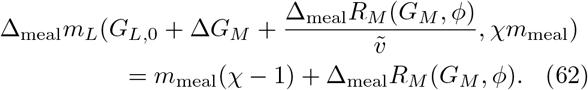

First we insist on robustness to *χ*, meaning that the muscle should achieve its target *ϕ* without knowing the composition of the meal. Since the muscle infers *m*_meal_ from its local glucose, this is only possible if Δ*G*_*M*_ (*m*_meal_, *χ*) is independent of *χ*. Differentiating Eq. (62) with respect to *χ* at fixed *m*_meal_ gives:

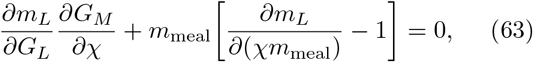

and therefore to have ∂*G*_*M*_ */*∂*χ* = 0,

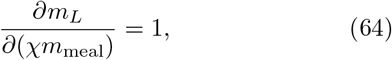

and thus

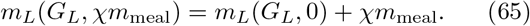

Notice that, with this condition satisfied, Eqs. (57)–(59) become *independent* of *χ*. Thus, when we insist on this robustness to meal composition, we can derive bounds by considering the case *χ* = 0, and they apply for all *χ*.

Our main equation thus becomes

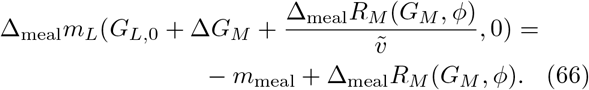

Now insist that Δ_meal_*R*_*M*_ (*G*_*M*_ (*ϕ, m*_meal_), *ϕ*) = *ϕm*_meal_. This gives the condition

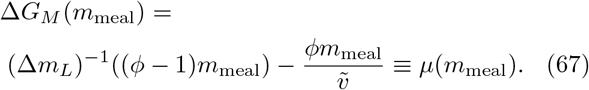

The muscle can invert this, controlling *ϕ* by setting

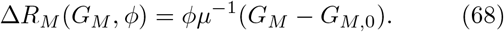

This is possible if, and only if, *µ*(*m*_meal_) is invertible in the relevant range. This requires that it is *monotonic*. In particular, it should be monotonically *increasing* : *R*_*M*_ (*G*_*M*_) must be an increasing function (glucose uptake should be higher at higher glucose), and thus we need *µ*(*m*_meal_) to be monotonically increasing rather than monotonically decreasing. Since 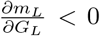, this means that we need

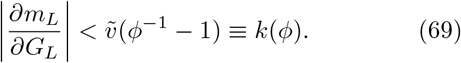

Since the liver cannot respond too aggressively to changes in *G*_*L*_, this places limits on the ability to prevent hyperglycemia in a meal. The precise values of these limits depend strongly on the assumed “operating range” over which the muscle can control *ϕ* (both the values of *ϕ* which can be achieved, and the values of *m*_meal_ for which they can be achieved): once this range is assumed, we can integrate Eq. (69) to obtain limits on *m*_*L*_(*G*_*L*_, *χm*_meal_) itself.

Concretely: assume that the body must allow all *ϕ < ϕ*_max_ for all *m*_meal_ *< m*_meal,max_. As we lower *ϕ* at fixed *m*_meal_, the required Δ_meal_*m*_*L*_ = (*ϕ* − 1)*m*_meal_ becomes more negative, and *G*_*L*_ thus rises. Thus, we may obtain the lower bound on the inverse function of *m*_*L*_ by first requiring 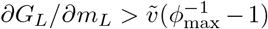 for −Δ_meal_*m*_*L*_ ≤ (1 − *ϕ*_max_)*m*_meal,max_ ≡ℳ, and then integrating the equation over *ϕ < ϕ*_max_, i.e.,

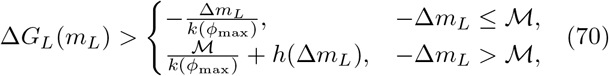

where

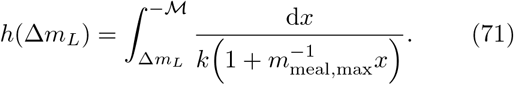

Evaluating the integral gives

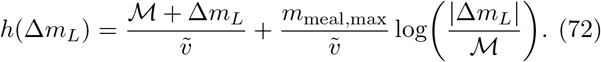

Once we have a lower bound on *G*_*L*_(*m*_*L*_, *χm*_meal_), we can turn this into a lower bound on *G*_*H*_ (*ϕ*) for a given *m*_meal_. The *χm*_meal_ terms again cancel, and we need

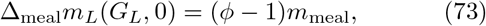

which gives us a lower bound on Δ_meal_*G*_*L*_. The gradient between *G*_*L*_ and *G*_*H*_ changes by an amount Δ_meal_(*m*_*U*_ + *m*_*L*_)*/v*_*LH*_ . These flux changes must balance Δ*R*_*M*_, and thus Δ_meal_*G*_*H*_ is smaller by *ϕm*_meal_*/v*_*LH*_, i.e.,

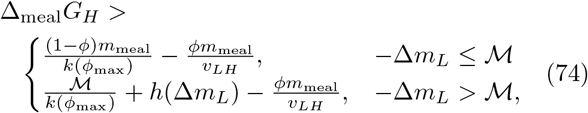

Equivalent equations appear in the main text.

### S6. CHANGES IN BLOOD FLOW

Here we derive Eq. (18) of the main text, and the trade-off it implies with the response to fluctuations in demand.

Under a change in *v*_*LH*_, we have

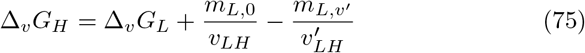

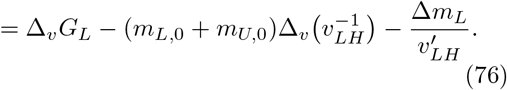

Now define *k*_*L*,eff_ as −Δ*m*_*L*_*/*Δ*G*_*L*_, giving:

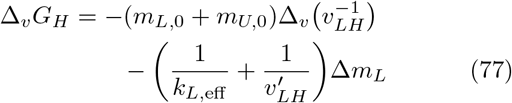

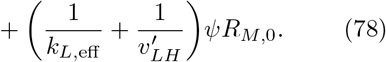

What happens under a fluctuation in demand? As stated in the main text, Eq. (19), we have

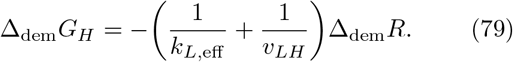

These two scenarios, a change in liver blood flow and a fluctuation in demand, define 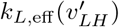 and *k*_*L*,eff_ (Δ_dem_*R*). Of course, both are really functions of the resulting *m*_*L*_. Thus, if we pick Δ_dem_*R* = −*ψR*_*M*,0_, resulting in the same total glucose demand in the two cases, then the two equations will involve the same *k*_*L*,eff_ and the two objectives will be linked, i.e.,

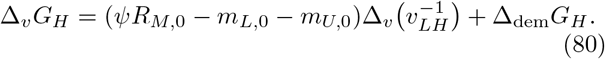

What if we want to put a bound on the trade-off between Δ_dem_*G*_*H*_ and Δ_*v*_*G*_*H*_ for Δ_dem_*R*≠ −*ψR*_*M*,0_? If we have a decrease in demand which is *larger* than *ψR*_*M*,0_, i.e., Δ_dem_*R <* − *ψR*_*M*,0_, then the best-case scenario for the resulting hyperglycemia is if the liver response has infinite slope beyond −*ψR*_*M*,0_, giving

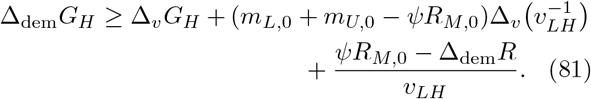

These two cases together give the plots in Fig. 4.

### S7. DETAILED CALCULATIONS OF TRADE-OFF ELIMINATION BY INSULIN

Here we show in detail how a circulating hormone eliminates the trade-offs imposed by local hormoneless control, as reported in the main text. As in the main text, insulin is produced in the pancreas at rate *βf* (*G*_*P*_), is cleared in the liver at rate *γ*_*L*_*I*_*L*_, and controls glucose production *m*_*L*_(*I*_*L*_). The pancreatic glucose *G*_*P*_ (*t*) follows the arterial concentration with a delay of a few seconds, Eq. (22), so the secretion rate responds to the recent history of *G*_*H*_ (*t*).

As a result, insulin in target tissues also reflects the history of *G*_*H*_ (*t*). The insulin component of the delay differential equations (DDEs) can be written in the form

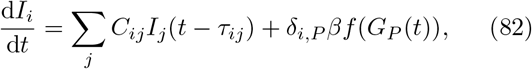

where *δ*_*i,j*_ is the Kronecker delta. As is standard, we define *U*_*ij*_(*t*) as the fundamental solution, i.e., the solution to the problem

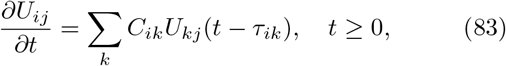

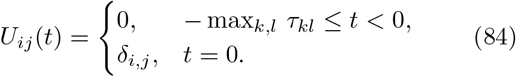

From the standard variation-of-constants formula [26], in the limit of long time, when transients die out, the solution of the inhomogeneous problem is then

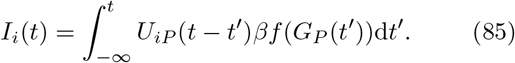

Thus, insulin in all tissues reflects the history of *G*_*H*_ (*t*).

In steady state, this simplifies immensely: we have

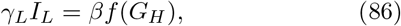

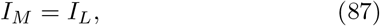

and thus, within our analyses of steady states, having a hormone secreted by the pancreas is completely equivalent to allowing each target tissue to respond directly to *G*_*H*_ .

This removes the exercise trade-off of the main text. In our model of exercise, the gradient between brain and heart does not increase, and total glucose balance, Eq. (42), gives

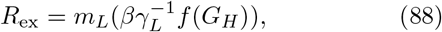

as stated in the main text. There is thus no longer any fundamental trade-off between fasting *G*_*H*_ and exercise *G*_*B*_.

The sensitivity to liver blood flow disappears as well, since the steady-state equations have no dependence on *v*_*LH*_ .

Finally, we consider meals. We allow liver and muscle to respond to insulin. As established above (Eqs. (85)–(87)), this is equivalent to allowing them to respond directly to *G*_*H*_, through functions 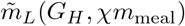 and 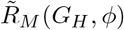.

It remains true that robustness to *χ* requires 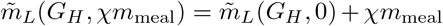, so it suffices to consider *χ* = 0.

The equivalent of Eq. (66) is

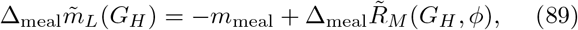

and again insisting that the muscle uptake change is *ϕm*_meal_, we obtain the condition

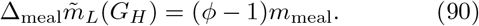

It is thus clear that, if the liver response 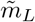 is known, the muscle can always control *ϕ*, by setting

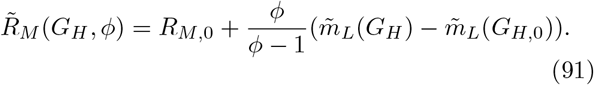

Thus, the trade-off shown in Fig. 3 is completely eliminated.

In reality, the muscle would implement this solution in terms of a response to insulin. With the simple model considered here, where insulin is cleared solely in liver, recall that we have *I*_*L*_ = *I*_*M*_, and thus we have the concrete implementation

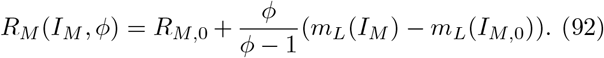

However, note that Eq. (85) still holds if (linear) insulin clearance takes place in other organs. Thus, although the formula will be more complicated, it is still possible to completely eliminate the tradeoff as long as *f* (*G*_*P*_) is monotonic.

### S8. SCHEMES WHICH TRY TO COMPENSATE FOR THE LIVER–GLUCOSE GRADIENT USING THE HISTORY OF LIVER GLUCOSE ARE UNSTABLE

The main text asserts that the estimator scheme of Eq. (21), in which the liver infers *G*_*H*_ from its own glucose and production rate, has an unstable steady state. Here we establish this numerically for the full model with delays, and then analytically for a simplified version, which shows that the instability does not depend on our particular parameters.

**FIG. S1.**
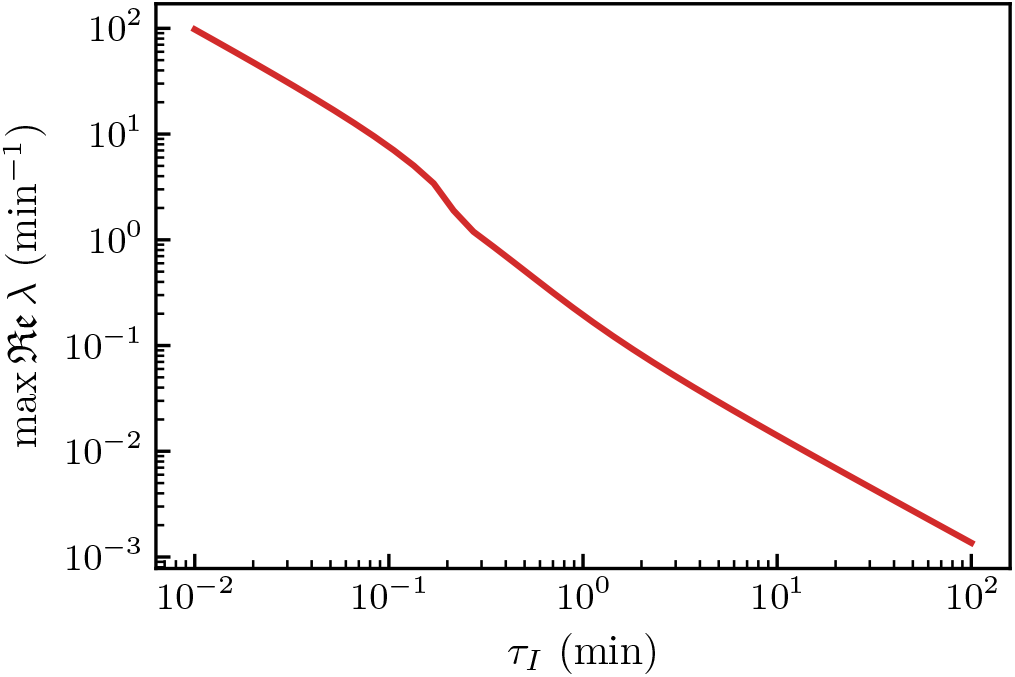
Largest real part of eigenvalues for linearized dynamics around steady state, with the scheme where *m*_*L*_ adjusts itself based on a local estimate of *G*_*H*_ (Eq. (21)). The real part of the stability-determining eigenvalue is always positive, indicating unstable dynamics.

We analyzed the steady states of the modified model that uses the history of liver glucose to estimate glucose in the heart, Eq. (21), using the DDE stability analysis code dde-biftool [27]. Linearization around a steady state produces a characteristic equation, which in a system with delays is transcendental and thus has infinitely many roots. We numerically approximate the roots with the most positive real part, which determine whether or not the steady state is linearly stable.

Linearizing Eq. (21) around a steady state (without loss of generality, assuming *u*^*′*^(0) = 1) yields

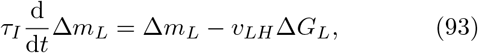

which we combine with the transport equations of our model (treating the uptake and production of the other organs as constant). We then study the stability-determining eigenvalue, max *Re λ*, as a function of *τ*_*I*_ . As shown in Fig. S1, the fixed point is unstable for any value of *τ*_*I*_ .

To see intuitively why this instability occurs, we simplify the analysis by (1) neglecting time delays and (2) grouping together the rest of the body (outside the liver). We thus introduce an associated volume *V*_body_ and glucose concentration *G*_body_. The linearized equations of motion around a steady state are then

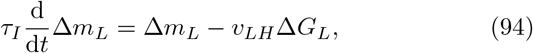

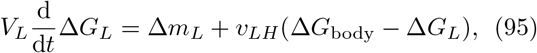

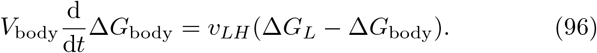

The linearized dynamics are of the form d**y***/*d*t* = **By**, for some matrix **B**, and **y** = (Δ*m*_*L*_*/v*_*LH*_, Δ*G*_*L*_, Δ*G*_body_). The characteristic equation of **B** is a cubic

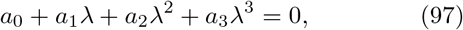

with

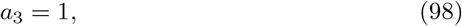

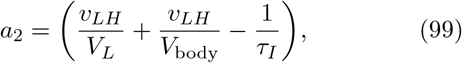

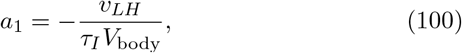

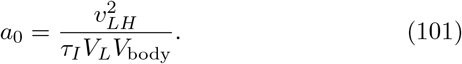

As long as all parameters are positive, *a*_1_ *<* 0 while *a*_0_, *a*_3_ *>* 0. The polynomial thus fails the Routh–Hurwitz stability criteria [28] and at least one of its roots has positive real part. Thus, at least one eigenvalue of **B** has positive real part, and the fixed points of the system Eqs. (94)–(96) are unstable. This suggests that the instability of the control scheme given by Eq. (21) is not sensitive to the specific parameters of our model.

From the simplified dynamics, we see that instability arises because trying to compensate for the liver–heart glucose difference requires including a term in d*m*_*L*_*/*d*t* which is proportional to *m*_*L*_(*t*) with a positive sign; we thus expect it to be a generic failure of schemes which try to estimate *G*_*H*_ using local liver information.

### S9. DELAYS DO NOT INTRODUCE INSTABILITY IN THIS MODEL, DUE TO WELL-MIXED ORGAN POOLS

Time delays can produce instabilities with high-gain control [29]. The bounds we derive in the main text allow the liver and muscle responses to be arbitrarily stiff, and those bounds would be unreachable if stiff responses were unstable here. It turns out that this does not happen in the simple model we have introduced, as we will now show.

We analyzed the steady states of our model using the DDE stability analysis code dde-biftool [27], as described in Section S8. In the linearized analysis, only the local gains (e.g., 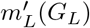 and 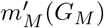) enter, and thus we do not need to assume any specific functional forms for *m*_*L*_ and *m*_*M*_ . For our model parameters, we plot max *Re λ* against these two gains, in Fig. S2. We see that increasing gain only decreases max *Re λ*. It thus appears that this model does not suffer from instability at large gain.

This traces directly to the well-mixed assumption, since in Eq. (6) the local response **g**_*i*_ depends on **x**_*i*_(*t*) with no delay, each organ being a single pool. The loop in which an organ senses its own glucose and adjusts its own flux therefore contains no delay at all, and every *τ*_*ij*_ sits instead in the transport terms between organs. Instability from delay requires a lag between measuring *G*_*L*_ and updating it through *m*_*L*_, which our model does not have. We thus expect that a model with more realistic internal structure within organs could still result in instabilities with highgain control.

**FIG. S2.**
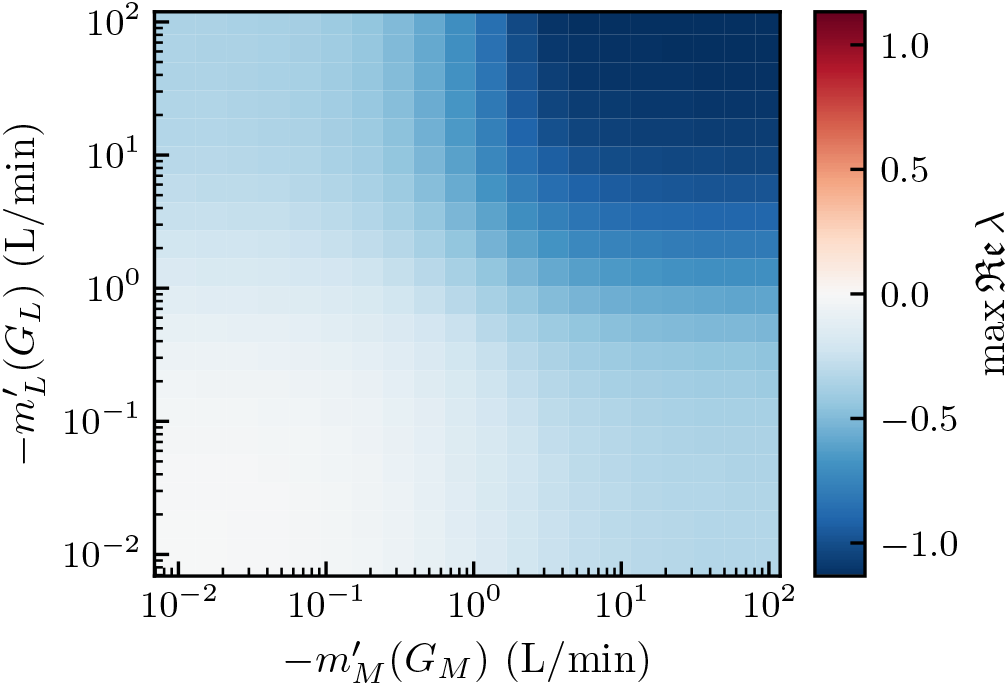
Largest real part of eigenvalues for linearized dynamics around steady state. The stability-determining eigenvalue always has negative real part, indicating stability, despite the presence of some time delays. The real part only becomes more negative for large gains, suggesting that instability does not develop even for arbitrarily large gains.

## Notes

### Competing Interest Statement

In the last 36 months, P.V. received consulting fees from Eli Lilly and Sanofi.

